# Quantifying motor protein copy number in super-resolution using an imaging invariant calibration

**DOI:** 10.1101/454900

**Authors:** Francesca Cella Zanacchi, Carlo Manzo, Raffaella Magrassi, Nathan D. Derr, Melike Lakadamyali

## Abstract

Motor proteins are nanoscale machines that convert the energy of ATP hydrolysis into the mechanical motion of walking along cytoskeletal filaments. In doing so, they transport organelles and help maintain sub-cellular organization. We previously developed a DNA origami-based calibration approach to extract protein copy number from super-resolution images. Using this approach, we show here that the retrograde motor protein dynein is mostly present as a single motor in the cytosol, whereas a small population of dynein along the microtubule cytoskeleton forms higher-order multimers organized into nano-sized domains. We further demonstrate, using dynein as a test sample, that the DNA origami-based calibration data we previously generated can be extended to super-resolution images taken under different experimental conditions, enabling the quantification of any GFP-fused protein of interest. Our results have implications for motor coordination during intracellular trafficking as well as for using super-resolution as a quantitative method to determine protein copy number at the nanoscale level.

## Introduction

Intracellular trafficking is an important biological process that facilitates the maintenance of spatial organization of organelles. Cytoplasmic dynein is the main motor protein responsible for intracellular retrograde transport. While the properties of single dynein motors have been extensively studied in vitro, the mechanisms regulating multiple motors that move organelles inside crowded cells are just starting to be uncovered^1^. Several experiments including optical trapping, fluorescence single step photobleaching and dark field microscopy, have shown that more than one dynein motor transports organelles in cells^2^^-^^8^. However, the molecular arrangement of these multi motor teams on organelles has not been directly visualized. This arrangement is an important determinant of how motor teams may cooperate or engage in a tug-of-war during transport^4^^,^^5^^,^^9^. It has also been suggested to play an important role in organelle motility, maturation and biogenesis^8^^,^^10^^-^^12^. Immuno-electron microscopy studies suggested that motor proteins can group in clusters on organelles^13^. More recently, cryo-electron microscopy studies showed that endosomal adapter proteins can accommodate two dynein dimers that sit in close physical proximity on one adapter protein^14^. In vitro single molecule experiments showed that when two dynein dimers are linked via an adapter protein, they can move faster than a single dynein motor^14^. Further, dynein was shown to form micro-domains on late phagosome-like compartments, and the formation of micro-domains was important for phagosome maturation^11^. These experiments combined together suggest that motors such as dynein can form a clustered organization containing multiple motors and this organization can be an important regulator of intracellular trafficking and cellular function. Further, the copy number of dynein motors that are present inside the micro- or nano-domains on the cargo membranes can be a crucial parameter for fine tuning trafficking^10^. However, quantifying the copy number of motor proteins like dynein inside the cellular milieu at the nanoscale level is highly challenging given their small size and high concentration. Nanometer scale spatial resolution is essential to be able to resolve the molecular arrangement of motors and count their copy numbers on microtubules, in the cytosol and when attached to organelles.

Super-resolution microscopy methods such as stochastic optical reconstruction microscopy (STORM) overcome the diffraction limit and allow visualizing the sub-cellular distribution of proteins with nanoscale spatial resolution^15^. However, quantifying protein copy number in super-resolution images is highly challenging due to the complex photophysics of fluorophores and the fact that labeling stoichiometry is often not one-to-one, especially when antibodies are used^16^^-^^18^. Previously, sparse images of single antibodies non-specifically adsorbed to cells were used as a reference for quantifying protein copy number in super-resolution images^19^^,^^20^, however, this method does not account for the unknown labeling stoichiometry. To overcome this problem, we and others have focused on developing calibration standards that can be used in conjunction with super-resolution microscopy^21^^-^^23^. In particular, we recently developed a versatile approach that uses a well-defined DNA origami structure as a calibration standard for super-resolution microscopy^24^. In this approach, we functionalized the DNA-origami with a defined number of GFP proteins, labeled the GFP using antibodies and carried out super-resolution imaging. By quantifying the number of localizations per GFP protein in the STORM image, we built a calibration curve that related the number of localizations to the GFP copy number. We further developed a statistical method that relies on fitting the distribution of localizations with a calibration function and allows quantification of the percentage of various oligomeric species within nanodomains at high resolution.

Here, we used this approach to quantify dynein copy number with nanoscale spatial resolution in both the cytosol and along the microtubule cytoskeleton of HeLa cells stably expressing dynein intermediate chain (DIC) fused to GFP (HeLa IC74 cells)^25^. A fully assembled dynein motor in its functional form is a dimer, containing two copies of DIC^26^. Super-resolution images revealed that cytosolic nano-clusters of DIC contained a large proportion of monomeric DICs likely corresponding to subunits not incorporated into a fully assembled motor complex. In addition, dimeric DICs, likely corresponding to a fully assembled dynein motor complex, were also present in the cytosol. Interestingly, along microtubules, the proportion of monomeric DIC decreased substantially. In addition, a small population of multimeric dyneins consisting of 2 or more motor complexes was found to be clustered into nano-sized domains along microtubules. These studies constitute the proof-of-concept demonstration that motors like dynein can indeed exist as nano-sized clusters containing multiple motor complexes in the physiological context.

In addition to applying our calibration approach to demonstrate the organization of dynein motors at the molecular level inside cells, we further extended the utility of our method. When we developed the DNA-origami calibration approach, we speculated that in order to use the calibration data we generated, the imaging of the protein of interest had to be carried out using the exact same imaging conditions as the one corresponding to the calibration experiment. Here, using dynein as a known, reference test sample, we further show that it is possible to identify proper calibration functions suitable for different imaging conditions. With this approach we can extend the calibration data from one set of experiments to a new set of experiments carried out under different experimental conditions, hence extending its versatility.

## Materials and Methods

### Sample preparation for super-resolution microscopy

HeLa IC74-mfGFP stably transfected cell line (from Takashi Murayama lab, Department of Pharmacology, Juntendo University School of Medicine, Tokyo, Japan) were plated on 8-well Lab-tek 1 coverglass chamber (Nunc) and grown under standard conditions (DMEM, high glucose, pyruvate (Invitrogen 41966052) supplemented with 10% FBS, 2 mM glutamine and selected with 400 µg/mL Hygromycin).

Cells were fixed with PFA (3% in PBS) at RT for 7’. Cells were then incubated at RT with blocking buffer (3% (wt/vol) BSA (Sigma) in PBS and 0.2% Tryton. In HeLa IC74-mfGFP stably transfected cells, dynein intermediate chain-green fluorescent protein (GFP) was immunostained with primary antibody (chicken polyclonal anti GFP, Abcam 13970) diluted 1:2000 in blocking buffer for 45 minutes at room temperature. Cells were rinsed 3 times in blocking buffer and incubated for 45 minutes secondary antibodies donkey-anti chicken labeled with photoactivatable dye pairs for STORM (Alexa Fluor 405-Alexa Fluor 647).

Human osteosarcoma U2OS cells (from ATCC, #HTB-96) were plated LabTek chambered coverglass (Nunc) and grown under standard conditions (DMEM, high glucose, pyruvate (Invitrogen 41966052) supplemented with 10% FBS. For GFP-tagged Nup133, cells were transfected using Fugene (FUGENE HD Transfection Reagent, Roche 04709705001) with the constructs (plasmid from Jan Ellenberg, EMBL, Heidelberg, pmEGFP-Nup133-s31401res, Euroscarf plasmid ref. P30728). Incorporation into the pore of the GFP-tagged Nup was facilitated by depletion of the endogenous protein performed by RNA interference, transfecting after 24h the cells with a matching siRNA (Nup133 SiRNA s31401, Thermo Fisher, Silencer Select siRNA s31401) and 3picomol of siRNA per well was used. After 60h cells were rinsed fixed with PFA (3%) for 7’.

### STORM microscopy

Imaging was performed with an oil immersion objective (Nikon, CFI Apo TIRF 100x, NA 1.49, Oil), repeated cycles of activation (405 nm laser), and readout (647 nm laser line) using TIRF illumination. During experiments the focus was locked through the Perfect Focus System (Nikon) and imaging was performed on an EMCCD camera (Andor iXon X3 DU-897, Andor Technologies).

Imaging condition 1: A custom-built total internal reflection fluorescence (TIRF) set-up built on an inverted Nikon Eclipse Ti microscope (Nikon Instruments) was used to acquire 85,000 frames at 25Hz frame rate. An excitation intensity of ~1KW/cm^2^ for the 647 nm readout (500 mW MPB Communications, Canada) and an activation intensity of ~ 25W/cm^2^ (100 mW, OBIS Coherent, CA) were used. Fluorescence emitted signal was spectrally filtered by a Quad Band filter (ZT405/488/561/647rpc-UF2, Chroma Technology) and selected by an emission filter (ZET405/488/561/647m-TRF, Chroma).

Imaging condition 2: A commercial N-STORM microscope (Nikon Intruments) was used to acquire 40,000 frames at 33Hz frame rate. An excitation intensity of ~0.9KW/cm^2^ for the 647 nm readout (300 mW MPB Communications, Canada) and an activation intensity of ~ 35W/cm^2^ (100 mW, Cube Coherent, CA) were used.

STORM imaging buffer was used containing GLOX solution as oxygen scavenging system (40 mg/mL^-1^ Catalase [Sigma], 0.5 mg/ml^-1^ glucose oxidase, 10% Glucose in PBS) and MEA 10 mM (Cysteamine MEA [SigmaAldrich, #30070-50G] in 360mM Tris-HCl).

### STORM Data analysis

Localization and reconstruction of STORM images were performed using custom software (Insight3, kindly provided by Bo Huang, University of California) by Gaussian fitting of the single molecules images to obtain the localization coordinates. The final image is obtained plotting each identified molecule as a Gaussian spot with a width corresponding to the localization precision (10nm) and corrected for drift.

A custom-written code written in Matlab^11^, implementing a distance-based clustering algorithm, was used to identify spatial clusters of localizations. The localizations list was first binned to 20 nm pixel size images that were filtered with a square kernel (7 x 7 pixel^2^) and thresholded to obtain a binary image. Only the localizations lying on adjacent (6-connected neighbours) nonzero pixels of the binary image were considered for further analysis. To select the sparse dynein contribution large clusters were filtered out setting a threshold on the maximum number of localizations (1000 localizations/cluster).

To estimate dynein copy number, we then used the DNA-origami-based method previously published^8^ and the procedure briefly described below. The quantitative approach uses DNA origami as a calibration standard and allows the copy-number estimation of a given protein from a general distribution of number of localizations by fitting the distribution to a linear combination of “calibration” functions.

Due to its stochastic nature, the STORM imaging procedure can be described as the mapping of a molecule into a random variable consisting of the number of localization N_1_ through a probability distribution *f*_1_. The shape of this distribution function can be determined by calibration experiments, such as imaging a large number of sparse molecules and counting the associated number of localizations per molecule.

This reasoning can be extended to protein clusters composed by a larger number of molecules. With two molecules A and B located at a distance shorter than the localization precision, the imaging procedure will provide, for each molecule, a random number of localizations N_1A_ and N_1B_ drawn from the distribution *f*_1_. Since the two molecules cannot be spatially resolved, it is impossible to associate the localizations to the different molecules. Therefore, we associate a number of localization N_2_ = N_1A_ + N_1B_ to the dimer. The distribution resulting from repeating this procedure over a large number of dimers will be the one associated to the sum of 2 identical random variables

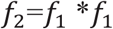

where the symbol * represents the convolution. The same reasoning can be iteratively extended to larger oligomers, obtaining that *f_N_* = *f_N_*_−1_ * *f*_1_.

The shape of the distribution for dimers and larger oligomers can be determined through a calibration experiment^24^.

If we perform an experiment aimed at imaging a given protein forming a heterogeneous population of oligomeric species, from the procedure described above we will obtain a distribution of number of localizations per cluster given by:

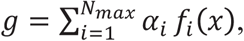

where *α_i_* are the fractions of i-mers and 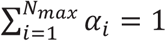. The fit of the experimental distribution with known functions *f_i_* gives the values of the coefficients *α_i_*, providing the copy-number estimation of a given protein^24^.

In principle, different data acquisition conditions provide different functions *f_i_* even if the same molecule is imaged. If we consider two microscopes (*M1* and *M2*) with similar localization precisions, the systems can be calibrated separately to obtain two different sets of calibration functions *f_i,M1_* and *f_i,M2_*. Alternatively, if only one of the two systems has been calibrated (e.g. *M1*), by imaging a simple, standard reference sample on both setups, we can obtain the calibration functions of the other setup (e.g. *M2*).

The imaging of the same reference sample in the different setups will produce

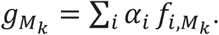

where 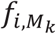 is a set of microscope-dependent calibration functions. If the functional form of 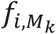 is known (log-normal in this case), the fitting of experimental data obtained from the same sample imaged with different setups must produce the same *α_i_* within the experimental and statistical uncertainty. Therefore, in the case we only have the calibration for one of the setups, imaging the same standard reference sample can be used to obtain the calibration functions for the other setup. Once the calibration function for the new imaging conditions is obtained by imposing the same output values for the coefficients *α_i_*, this new calibration function can be used to fit new data acquired using the second setup.

To implement this approach, we calculated the coefficients *α_i_* for dynein intermediate chain (DIC) imaged with microscope 1 (*M1*) using the calibration functions obtained for *M1* through the DNA origami approach. Since a fully assembled dynein motor complex is homodimeric, containing 2 copies of DIC, in this procedure we included all the species corresponding to even multiples of DIC, while also considering the existence of a flexible population of monomeric DIC subunits. For DIC imaged on microscope 2 (*M2*) with different acquisition settings, fitting to a linear combination of calibration functions was performed by imposing the coefficients *α_i_* corresponding to fully assembled motor complexes to be the same as those determined on M1 within a 10% uncertainty. We run the fit on the free parameters µ and σ and thus obtain the lognormal calibration function for the new microscope.

## Results

### A small population of dynein is clustered into nano-sized domains composed of multiple dynein motors along microtubules

To determine the nanoscale distribution and copy number of dynein, we used HeLa cells stably expressing dynein intermediate chain (DIC) fused to GFP (HeLa IC74). The super-resolution images of dynein labeled with anti-GFP primary and secondary antibodies revealed nano-sized clusters (**Figure 1**). These clusters were similar whether cells were fixed using PFA or methanol-ethanol (**Figure S1**, Methods), suggesting that dynein organization is independent of the fixation method. We segmented these clusters using a distance-based cluster identification algorithm that we previously developed^27^. Overlay of the super-resolution images with wide-field images of microtubules showed that some of these clusters overlapped with the microtubule cytoskeleton (**Figure 1a**) whereas others were distributed in the space in between microtubules and were most likely cytosolic (**Figure 1b**). Using the wide-field image of microtubules as a mask, we manually segmented and classified sub-regions in the dynein super-resolution image into two categories: overlapping with microtubules (**Figure 1a and inset**) and cytosolic (**Figure 1b and inset**). Since a diffraction limited image of microtubules was used for this analysis and both the wide-field and super-resolution images are 2D projections, the population of dynein overlapping with microtubules likely contains a mixed population of microtubule associated and cytosolic dynein motors. The cytosolic population, on the other hand, should consist of motors predominantly not associated to microtubules.

**Figure 1:**
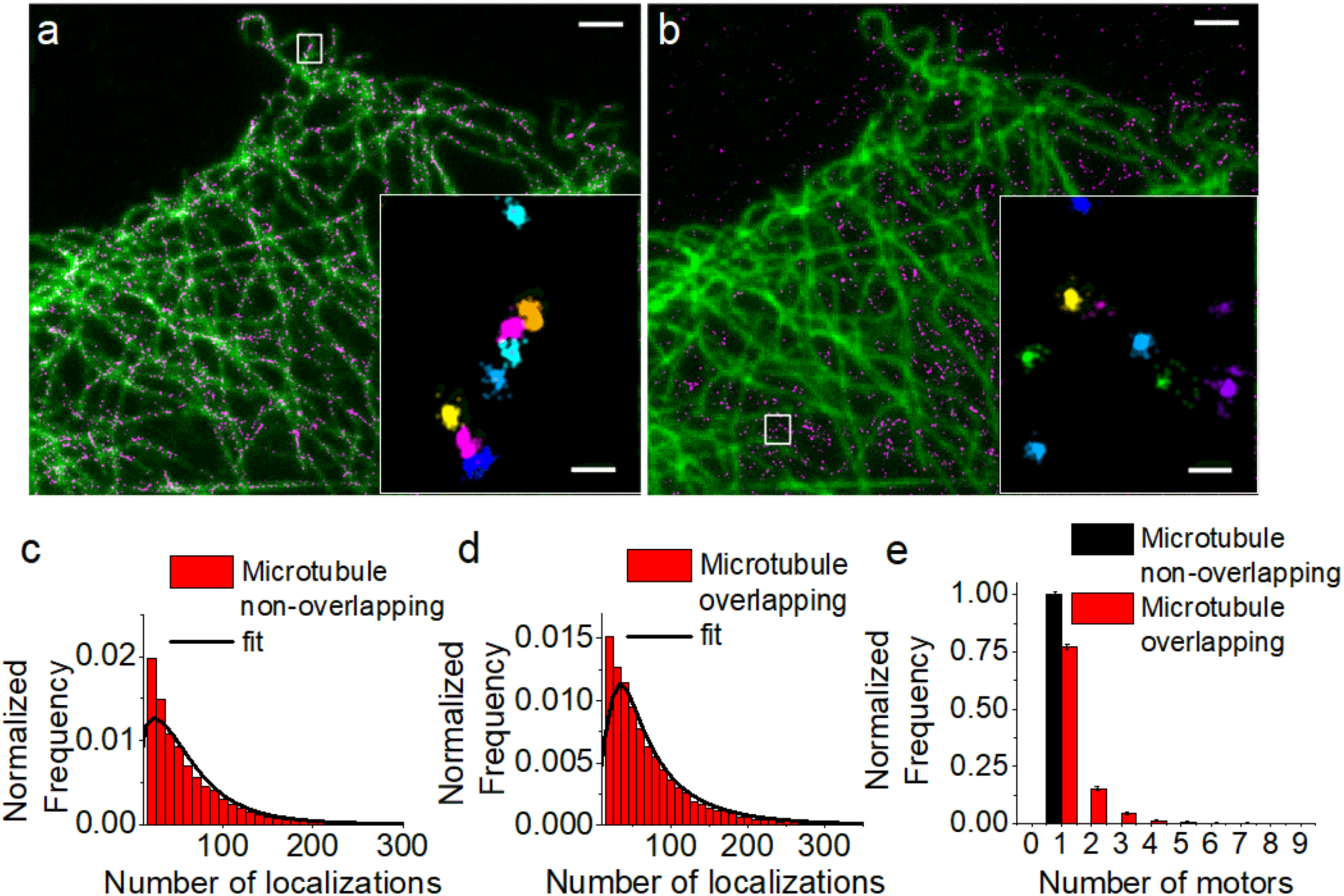
Super-resolution imaging and quantification of dynein organization in cells reveals the presence of dynein multimers organized into nanoclusters along microtubules: (**a,b**) STORM imaging of dynein intermediate chain fused to GFP and immunostained with anti-GFP primary and AlexaFluor405/AlexaFluor647 labelled secondary antibodies (magenta). Wide-field images of microtubules (green) was used to separate dynein overlapping with the microtubule cytoskeleton (**a**) (n=6 independent experiments, n=6958 clusters) from cytosolic dynein not associated to microtubules (**b**) (n=6 independent experiments, n=6345 clusters). Zooms of the regions inside white boxes are shown as insets after cluster analysis to segment the individual dynein nanoclusters. Scale bars: 2µm (**a, b**), 200nm (**insets**). Distributions showing the number of localizations per dynein nanocluster non-overlapped **(c)** and overlapped with microtubules (**d**). The distributions were fit to a linear combination of calibration functions (black lines). The goodness of fit was evaluated with a reduced-χ^2^ test (see **Figure S3a, b**). (**e**) Copy-number for dynein overlapped with (red) and not-overlapped with (black) microtubules obtained from the fit and normalized to show the homodimeric, fully assembled motor complexes only. The maximum number of motors was calculated minimizing the objective function both for dynein overlapped (**Figure S2a**) and non-overlapped (**Figure S2b**) with microtubules.

To quantify the copy number of dynein motors in the clusters, we used the calibration data that we previously generated^24^. Dynein is a homodimer containing two copies of DIC. Our previous calibration data was obtained by using a purified, modified dimeric dynein motor in which GFP was fused to the motor’s cargo binding domain^24^. Therefore, we could determine a calibration distribution (*f_1,M1_*, in which *M1* refers to microscope acquisition setting 1) describing the probability distribution of the number of localizations for a fully assembled dynein motor having two copies of GFP^24^. We also inferred the calibration function for a monomer containing only one copy of GFP (*f_0.5,M1_*). Since a proportion of the DIC may not be incorporated into a fully assembled, functional dynein motor complex and may exist as a monomer, we fit the number of localizations per cluster to a linear combination of the calibration distributions *f_i,M2_* (**Figure 1c and d**) with values *i*=0.5, 1, 2, 3…. In this case, a copy number of 0.5 would correspond to monomeric DIC, a copy number of 1 would correspond to a homodimeric DIC representing a fully assembled dynein motor complex, a copy number of 2 would correspond to 2 fully assembled dynein motor complexes etc… The fitting was stopped at some *i*=N_max_ for which the objective function was minimized (**Figure S2**) and the goodness of fit was evaluated using the reduced-χ^2^ value (**Figure S3a and Figure S3b**). This quantification revealed that the cytosolic dynein clusters were a combination of monomeric DIC (likely not incorporated into a motor complex) and homodimeric DIC (single copy of dynein motor complex) (**Figure S4a**). The monomeric population was dramatically reduced in the population of dynein clusters overlapping with microtubules (**Figure S4b**). Furthermore, this population also contained multimeric dynein motor complexes consisting of 2 or more dyneins inside the nanocluster **(Figure 1e, red bars)**, which was missing from the cytosolic population (**Figure1e, black bars**). These results suggest that along the microtubules, DIC is predominantly incorporated into a homodimeric full motor protein complex unlike the cytosolic population and these full motor complexes can further cluster together into nano-sized domains forming higher order dynein multimers (**Figure 2e**). To further evaluate the validity of the colocalization analysis in which wide-field images of microtubules were used as masks, we determined the copy number of dynein from regions that were randomly selected. We reasoned that these randomly selected regions should resemble the cytosolic population since microtubules occupy a minor proportion of the cytosol. Indeed, the copy number of dynein in nanoclusters from randomly selected regions was identical to the cytosolic nanoclusters (**Figure S4b and c**). While this analysis does not exclude that the nanoclusters overlapping with microtubules may be a mixed population of cytosolic and microtubule associated dynein, it validates that the differences between the cytosolic and the microtubule overlapping nanoclusters are indeed due to dynein associated to microtubules. We are likely underestimating the actual difference in these two populations of dynein nanoclusters.

**Figure 2:**
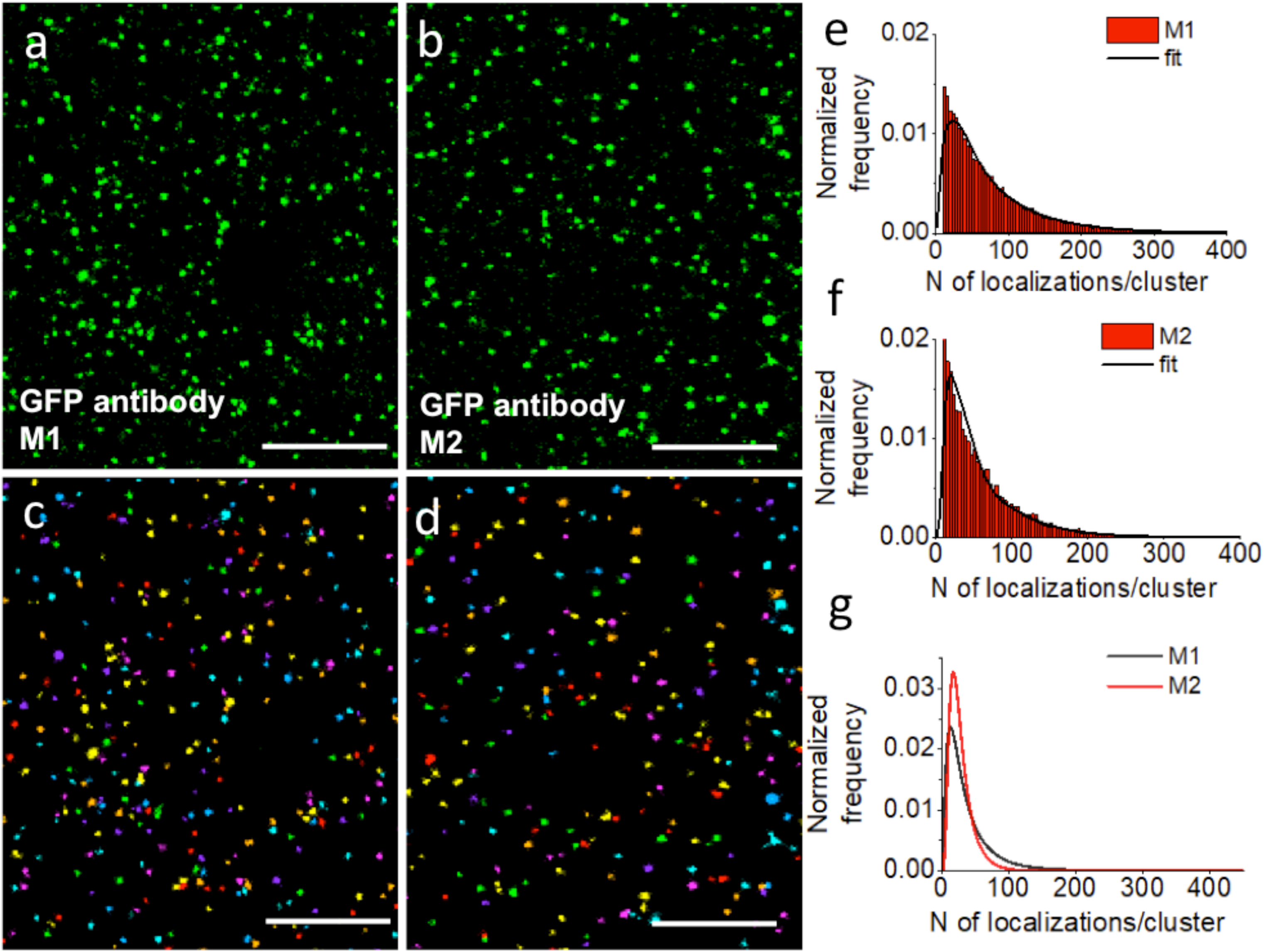
New calibration functions can be obtained for different imaging conditions using dynein quantification as a reference. Super-resolution images of dynein intermediate chain fused to GFP and immunostained with anti-GFP primary and AlexaFluor405/AlexaFluor647 labelled secondary antibodies under different imaging conditions (**a,b**) and corresponding cluster analysis (**c,d**). Distributions showing the number of localizations quantified from nanoclusters in the entire cell and the corresponding fit to linear combination of calibration functions for different imaging conditions (**e,f**). The goodness of fit was evaluated with a reduced-χ^2^ test (see **Figure S3c and d**). Calibration functions for the old (M1) (**g, black**) and for the new (M2) imaging conditions (**g, red**). Scale bars 2 µm.

### Calibration functions can be obtained for different imaging conditions by using a reference sample

In order to quantify the copy number of dynein motor protein complexes in the intracellar milieu, the super-resolution images were taken using the exact same imaging conditions (i.e. *M1*) that were used to obtain the DNA-origami calibration data. To determine the impact of image acquisition settings on the quantification, we recorded a new set of super-resolution images of dynein by once again labeling DIC with anti-GFP antibodies and modifying the acquisition settings (*M2,* see Methods) (**Figure 2a-d**). The modified image acquisition parameters indeed changed the distribution of the number of localizations per cluster extracted from the super-resolution images (**Figure 2e, f**). Therefore, fitting this new set of experimental data (**Figure 2f**) with the previous calibration function (obtained for *M1*) would produce inaccurate results for the copy number distribution of dynein within the nanoclusters. We expect that a change in the imaging protocol should also produce a different calibration function, whereas, imaging equivalent “reference” samples should produce the same copy number distribution with the same coefficients *α_i_* (i.e. the proportion of each i-meric species should remain the same). The latter assumption can be used to obtain the calibration function for the new imaging condition (**Figure 2g, red**) by imposing similar values for *α_i_* within the experimental uncertainty as constraint and extracting the parameters for the calibration function (i.e. µ and σ of the log-normal distribution) from the fit. The number of localizations obtained for single dynein clusters in HeLa cells fit well to a linear combination of calibration functions both under the old (*M1,* **Figure 2e**) and new imaging conditions (*M2,* **Figure 2f**) as also shown by the reduced-*χ*^2^ values for the goodness of fit (**Figure S3c, d**). By studying the curvature of the loglikelihood function around the estimator values, we could determine the standard error of the free parameters, showing that they are well constrained (mu= 3.227 ± 0.007, sigma=0.569 ± 0.007). The match between the copy number distribution results obtained for the two different imaging conditions was further checked *a posteriori* by a *χ*^2^ test providing a p-value≃1, thus excluding significant differences between the two sets (**Figure S5**). This result is expected since the two copy number distributions were imposed to be the same within experimental uncertainty.

To validate the method, we extracted dynein copy number using the new calibration function on a new set of images of DIC under the *M2* imaging conditions (**Figure 3a**). It’s important to note that these new images constituted an independent data set, with no overlap with the one used for extracting the *M2* calibration function. Further, since we did not obtain microtubule images in this case, the quantification was carried out for dynein nanoclusters in the entire cell and this analysis was repeated for the images obtained using *M1* for comparison. Although we considered the existence of a monomeric DIC population in the fit, we focused on determining and comparing the copy number distribution of fully assembled homodimeric dynein motor complexes since these are the more biologically relevant population. Once again, we obtained a large proportion of dynein in single copy and smaller proportion of multimeric dynein motor complexes (**Figure 3b, black bars**). The small differences in dynein copy numbers compared to the previous quantification using the *M1* imaging conditions (**Figure 3b, red bars**) is expected from the cell to cell variability in the dynein organization as well as experimental uncertainties. A *χ*^2^ test showed no significant difference between the two results (p-value≃1), further validating our approach.

**Figure 3:**
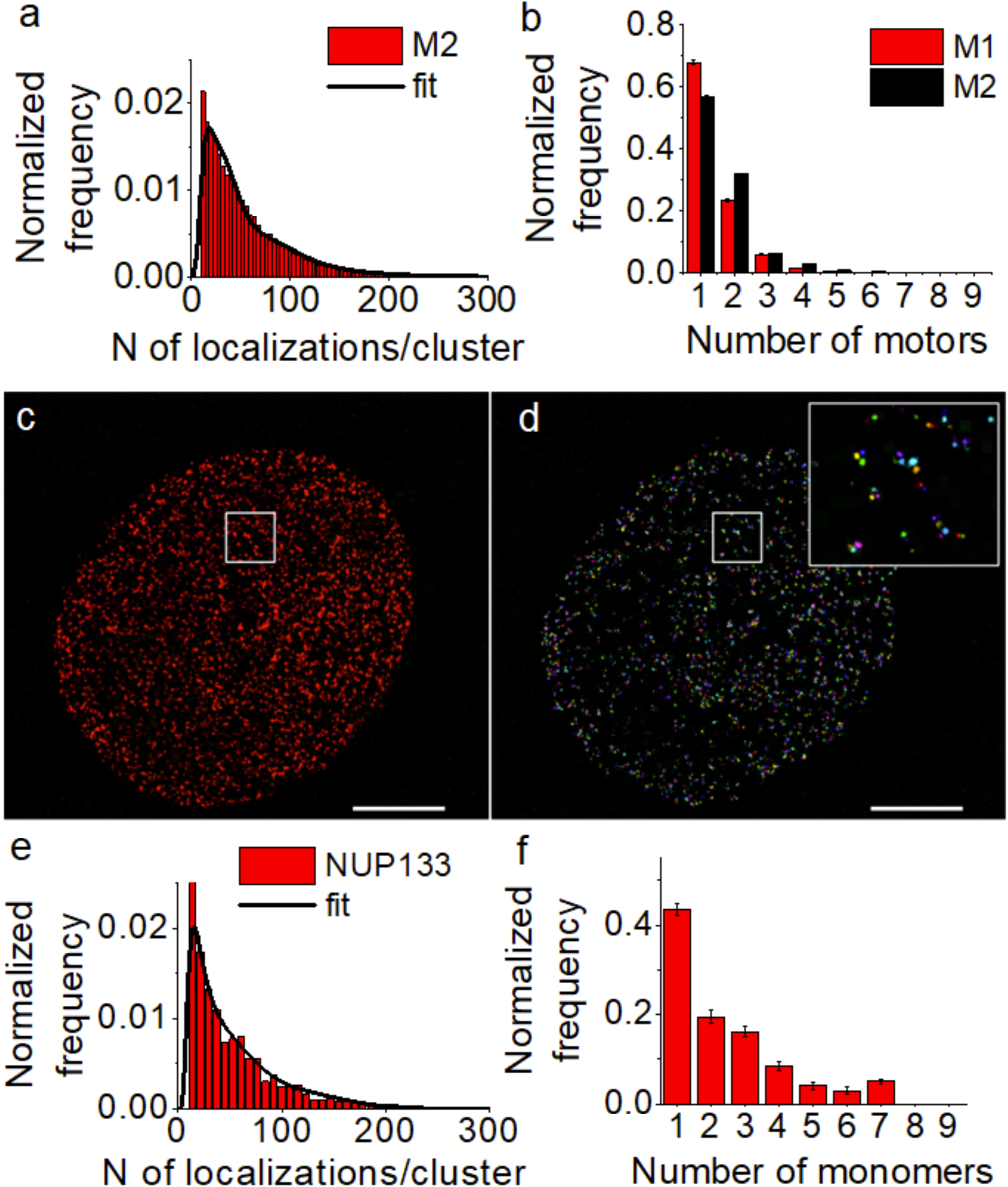
Validation of the new calibration function using dynein and nuclear pore complex. (**a**) Distribution showing the number of localizations per cluster obtained from super-resolution images of DIC that were not used for determining the new calibration function shown in Figure 2. The corresponding fit to a linear combination of the new (M2) calibration function is shown as a black line. The goodness of fit was evaluated with a reduced-χ^2^ test (see **Figure S3e**). (**b**) Dynein copy number distribution in the entire cell quantified using M1 (red) (n=7 independent experiments, n=25427 clusters) and M2 (black) calibration functions (n=3 independent experiments, n=18490 clusters). The copy numbers were normalized to show the homodimeric, fully assembled motor complexes. The match between the copy number distribution of dynein obtained under M1 and under M2 was determined by a χ^2^ test (χ^2^ value of 0.0313 provides a p-value=1). Super-resolution image (**c**) and cluster analysis (**d**) of NUP133 expressing GFP immunostained with AlexaFluor405/AlexaFluor647 labelled secondary antibodies and obtained using M2 imaging conditions. A zoom of the region inside the white box is shown (**d,inset**). **(e)** The number of localizations distribution for each cluster corresponding to one fold of the NPC and the corresponding fit to the linear combination of calibration functions for M2 imaging condition. The goodness of fit was evaluated with a reduced-χ^2^ test (see **Figure S3**f). (**d**) GFP copy-number distributions estimated for single clusters of Nup133 corresponding to one fold of the NPC. (N=1 experiment, total number of clusters analyzed n=4982).

To further test the performances of the method and validate its application, we quantified the copy number distribution of NUP133 (a subunit of the nuclear pore complex) fused to GFP. The subunit copy number of NUP133 has been extensively characterized by a number of methods^28^. The nuclear pore complex is a ring shaped structure with 8-fold symmetry. Each fold contains 4 copies of NUP133. Therefore, when over-expressing NUP133 fused to GFP, we expect each fold to contain between 1-4 copies of GFP along with endogenous, untagged NUP133. We immunostained the GFP in cells over-expressing NUP133-GFP, obtained super-resolution images using the new imaging conditions (*M2*) (**Figure 3c**) and performed cluster analysis (**Figure 3d**) to segment a single fold of the NPC (**Figure 3d, inset**). In order to determine the copy number of GFP in each NUP133 cluster, we fit the distribution of the number of localizations per cluster (**Figure 3e**) to a linear combination of the new calibration distributions (*f_i,M2_*). This quantification revealed that majority of the NUP clusters were indeed composed of 1-4 copies of GFP, as expected (**Figure 3f**).

## Discussion

Intracellular transport is a fundamental biological process regulated by many mechanisms including regulation through the microtubule cytoskeleton^29^, adapter and accessory proteins^1^, as well as motor-protein copy number and the molecular arrangement of motor-proteins on the vesicle membrane^10^^-^^12^^,^^14^. Indeed, previous modeling work predicted that the copy number and arrangement of motors are important parameters for regulating transport^10^. However, determining the precise molecular organization and copy number of motors in the cellular context has been difficult given the small size of sub-cellular compartments and motor-proteins as well as the crowded intracellular environment. Here, using super-resolution microscopy and a DNA origami-based calibration approach that we previously developed^24^, we show that dynein can cluster into nano-sized domains containing multiple dynein motor complexes, in particular along microtubules. Our results are consistent with the cryoEM data showing that endosomal adapter proteins can recruit two copies of dynein in very close spatial proximity to one another^14^. These nanoclusters were smaller than the dynein microdomains observed on purified phagosomes^11^. In the future, it would be exciting to determine whether these dynein clusters are associated to specific sub-cellular vesicles, whether kinesin family motors can also cluster, and how clustering of multiple-motors may impact transport inside the cell. Further, it would be exciting to determine the mechanisms that mediate clustering of motor proteins such as dynein. Clustering may be mediated by the cytoskeleton itself, by lipid composition of vesicles^11^ or by adapter and accessory proteins^14^. The methods we outline here enable these future mechanistic and functional studies.

We further show that in order to use the DNA origami-based calibration data we previously generated, it is not strictly necessary to follow the exact same imaging conditions as the calibration experiment or to repeat the calibration experiment using new imaging conditions. Instead, it is sufficient to image a reference sample under the two imaging conditions as a benchmarking standard. If such an experiment is performed, it is possible to extract a new calibration function by imposing that the reference sample imaged under different imaging conditions must contain the same copy number distribution. This new calibration function can then be applied to determine the copy number of any GFP-labeled protein of interest imaged using the new acquisition settings. We validated this approach by imaging and determining the copy number of both dynein and the NUP133 subunit of the nuclear pore complex imaged under different imaging conditions. Here, we used HeLa IC74 cells stably expressing dynein intermediate chain fused to GFP as the “reference sample” since we quantified the copy number of dynein complexes in this cell line. However, our approach is not limited to using this cell line as a reference and other, more standard reference samples, such as commercially available gene-edited GFP reporter cell lines can be used. We found that both µ and σ of the log-normal calibration distribution was well constrained by the fit. We expect that how well constrained these two parameters are will depend on the shape of the oligomeric distribution in the reference sample and hence this consideration should be taken into account when choosing an appropriate reference sample.

Our approach not only accounts for any differences in image acquisition settings but also differences in antibody labeling (e.g. the number of fluorophores per antibody) or the imaging buffer. Matching these experimental conditions among different laboratories can be challenging. Further, carrying out a calibration experiment using DNA origami requires extensive sample preparation including protein purification and DNA origami functionalization. Alternative calibration standards that take advantage of protein complexes with known copy numbers expressed in cells have also been developed^21^^,^^22^^,^^30^^,^^31^. The use of these calibration standards also require special care since expressing these protein complexes in cells such that they are fully assembled and matured without forming aggregates is challenging. Therefore, the approach we outline here, consisting of using a simple, readily available and easy to prepare reference standard to calibrate and match different acquisition settings will help make stoichiometry quantification more universal.

## Acknowledgements

HeLa IC74 cell line was a kind gift of Takashi Murayama, Department of Pharmacology, Juntendo University School of Medicine, Tokyo, Japan. We thank Dr. Angel Sandoval Alvarez for helping with cell maintenance and transfections. M.L. acknowledges funding from the Perelman School of Medicine, University of Pennsylvania, the Fundació Cellex Barcelona, European Union Seventh Framework Programme under the European Research Council grants 337191-MOTORS and Spanish Ministry of Economy and Competitiveness and the Fondo Europeo de Desarrollo Regional (FEDER) grant FIS2015-63550-R (MINECO/FEDER). F.C.Z. acknowledges funding from the “Severo Ochoa” Programme for Centres of Excellence in R&D (SEV-2015-0522). C.M. acknowledges funding from the Spanish Ministry of Economy and Competitiveness and the European Social Fund through the “Ramón y Cajal” program 2015 (grant RYC-2015-17896) and the “Programa Estatal de I+D+i Orientada a los Retos de la Sociedad” (grant BFU2017-85693-R).

